# An N-terminal delivery domain defines a new class of polymorphic T6SS effectors in *Enterobacterales*

**DOI:** 10.1101/2023.07.16.549128

**Authors:** Andrea Carobbi, Simone di Nepi, Eran Bosis, Dor Salomon, Guido Sessa

**Author notes:** To whom correspondence should be addressed: DS. We are deeply saddened to report the untimely loss of our friend and colleague Prof. Guido Sessa, who passed away on July 4, 2023.

## Abstract

The type VI secretion system (T6SS), a widespread protein delivery apparatus, plays a role in bacterial competition by delivering toxic effectors into neighboring cells. Identifying new T6SS effectors and deciphering the mechanism that governs their secretion remain major challenges. Here, we report two orphan, antibacterial T6SS effectors in the pathogen *Pantoea agglomerans* (*Pa*). These effectors share an N-terminal domain, PIX, that defines a widespread class of polymorphic T6SS effectors in *Enterobacterales*. We show that the PIX domain is necessary and sufficient for T6SS-mediated effector secretion and that PIX binds to a specialized *Pa* VgrG protein, outside of its C-terminal toxic domain. Our findings underline the importance of identifying and characterizing new delivery domains in polymorphic toxin classes as a tool to reveal novel effectors and shed light on effector delivery mechanisms.

## Introduction

Bacteria secrete toxic proteins to combat their rivals and gain a competitive advantage (Klein et al., 2020). Many Gram-negative bacteria employ the Type VI secretion system (T6SS), a contact-dependent protein delivery apparatus that transports toxic effectors directly into neighboring cells (Hood et al., 2010; Jana and Salomon, 2019; Mougous et al., 2006; Pukatzki et al., 2006). The effectors are loaded onto a missile-like structure comprising a tube of stacked hexameric Hcp rings capped by a spike complex made of a VgrG trimer and a sharpening PAAR repeat-containing protein (Cherrak et al., 2019). The contraction of an engulfing sheath structure propels the missile and effectors out of the cell (Basler et al., 2012; Cherrak et al., 2019).

T6SS effectors come in two flavors: (i) specialized effectors, which are the secreted structural proteins Hcp, VgrG, and PAAR carrying a C-terminal toxic domain extension; and (ii) cargo effectors, which are toxic proteins that non-covalently load onto a secreted structural T6SS component, often aided by an adaptor protein, a co-effector, or a tether protein (Dar et al., 2022; Jana and Salomon, 2019; Kanarek et al., 2022; Unterweger et al., 2017). Most effectors studied to date mediate antibacterial toxicity by targeting conserved cell components, such as: nucleic acids, the peptidoglycan, or the membrane (Klein et al., 2020); these antibacterial effectors are encoded together with a cognate immunity protein that protects against self- or kin-intoxication.

The lack of a canonical secretion signal in T6SS cargo effectors hinders their identification. Nevertheless, several strategies have successfully revealed T6SS cargo effectors, including comparative proteomics (Altindis et al., 2015; Ray et al., 2017; Russell et al., 2012; Salomon et al., 2014, 2015; Tchelet et al., 2023), transposon screens (Dong et al., 2013), and computational approaches, such as comparative genomics (Fridman et al., 2020), the investigation of genes encoding proteins of unknown function within the neighborhood of T6SS gene cluster or operons (Jana et al., 2019, 2022; Liang et al., 2015) and the presence of domains associated with secreted polymorphic toxins (Fridman et al., 2022; Kanarek et al., 2022; Salomon et al., 2014). Several N-terminal domains that define classes of polymorphic toxins and play a role in their delivery are associated with T6SS effectors: (i) MIX (Salomon et al., 2014), (ii) FIX (Jana et al., 2019), (iii) RIX (Kanarek et al., 2022), and (iv) Rhs repeats (Koskiniemi et al., 2013). In contrast to the first three, Rhs is not strictly associated with T6SS. Moreover, unlike MIX, FIX, and Rhs, which are widespread in Gram-negative bacterial families, RIX is restricted to *Vibrionaceae*. Notably, many cargo effectors do not have a defined N-terminal delivery domain, suggesting that additional polymorphic T6SS delivery domains remain to be revealed.

The genus *Pantoea* comprises ecologically diverse species, including the economically important plant pathogens *Pantoea stewartii* (Braun, 1982), which is the causal agent of a devastating disease in maize, and *Pantoea ananatis* (Coutinho and Venter, 2009), which causes disease on a broad range of host plants. *Pantoea agglomerans* (*Pa*) is an epiphytic and endophytic bacterium. It is found in diverse natural and agricultural habitats (Lindow and Brandl, 2003; Lorenzi et al., 2022). Although mainly considered beneficial to the environment (Dutkiewicz et al., 2016), *Pa* can rarely cause human diseases (Büyükcam et al., 2018; Cruz et al., 2007). Two phytopathogenic *Pa* pathovars induce plant tumors known as galls: *Pa* pv. *gypsophilae* (*Pag*) and *Pa* pv. *betae* (*Pab*). These pathovars inflict substantial losses in beet (*Beta vulgaris*) crops and prevent root development of gypsophila (*Gypsophila paniculata*) cuttings used in the flower industry (Burr, 1991; Cooksey, 1986).

We recently reported that the *Pab* T6SS is an active antibacterial apparatus that provides a competitive advantage over other plant colonizers, and we functionally characterized two effectors encoded within the T6SS gene cluster (Carobbi et al., 2022). We also proposed that additional antibacterial cargo and specialized effectors are found within rapidly evolving islands in the T6SS gene cluster (Carobbi et al., 2022). However, whether T6SS effectors are encoded outside the *Pab* T6SS gene cluster remained unknown. Here, we reveal two orphan, antibacterial T6SS effectors in *Pab*. We show that these effectors share a previously unknown N-terminal domain, PIX, which is necessary and sufficient for T6SS-mediated secretion via interaction with the VgrG component of the T6SS. Importantly, we find that PIX can be used as a marker to identify novel orphan T6SS effectors in *Enterobacterales*.

## Results

### Revealing orphan antibacterial T6SS effectors in *P. agglomerans* pv. *betae*

T6SS specialized effectors and T6SS cargo effectors may contain similar C-terminal toxic domains (Fridman et al., 2020; Jana et al., 2019; Salomon et al., 2014), suggesting that a protein containing homology to the C-terminus of a T6SS specialized effector may be a T6SS cargo effector. Therefore, when we found a *Pab* protein, WP_064704874.1, with homology to the C-terminus of a predicted PAAR domain-containing effector from the plant pathogen *Ralstonia*, we hypothesized that it is a T6SS cargo effector.

The C-terminus (amino acids 213-348) of WP_064704874.1, hereafter referred to as Pse3 (<u>P</u>antoea type <u>s</u>ix <u>e</u>ffector 3; following the previously reported nomenclature of *Pab* T6SS effectors (Carobbi et al., 2022)), shares 56% identity with the C-terminus (amino acids 250-385) of the PAAR domain-containing protein WP_011003305.1 (**Fig. 1A**). Interestingly, we identified two *Pab* proteins that are homologous to the N-terminus of Pse3: WP_022625183.1 and WP_231115755.1, hereafter referred to as Pse4 and Pse5, respectively (**Fig. 1A**). Pse4 contains a C-terminal domain predicted by the NCBI Conserved Domain Database (CDD) (Marchler-Bauer et al., 2007) to belong to the Colicin_D nuclease superfamily (Tomita et al., 2000). Its N-terminus (amino acids 1-215) shares 72% identity with the N-terminus of Pse3 (amino acids 1-213). <u>Pse5</u> is a shorter predicted protein; the protein encoded downstream is a predicted member of the HNH nuclease superfamily (WP_231115754.1; CDD) (Zhang et al., 2012). DNA sequence analysis indicated that a stop codon found after the codon for amino acid 157 of Pse5 (validated using Sanger sequencing) prevented it from forming a single open reading frame with the downstream gene. Notably, nearly identical proteins in which the stop codon is substituted by a tyrosine codon (e.g., WP_089414745.1) are encoded in other *Pantoea* strains; they share 58% identity with the N-terminus of Pse3 (**Supplementary Fig. S1**). Because of the apparent truncation of Pse5 in *Pab*, we excluded it from subsequent analyses.

**Figure 1.**
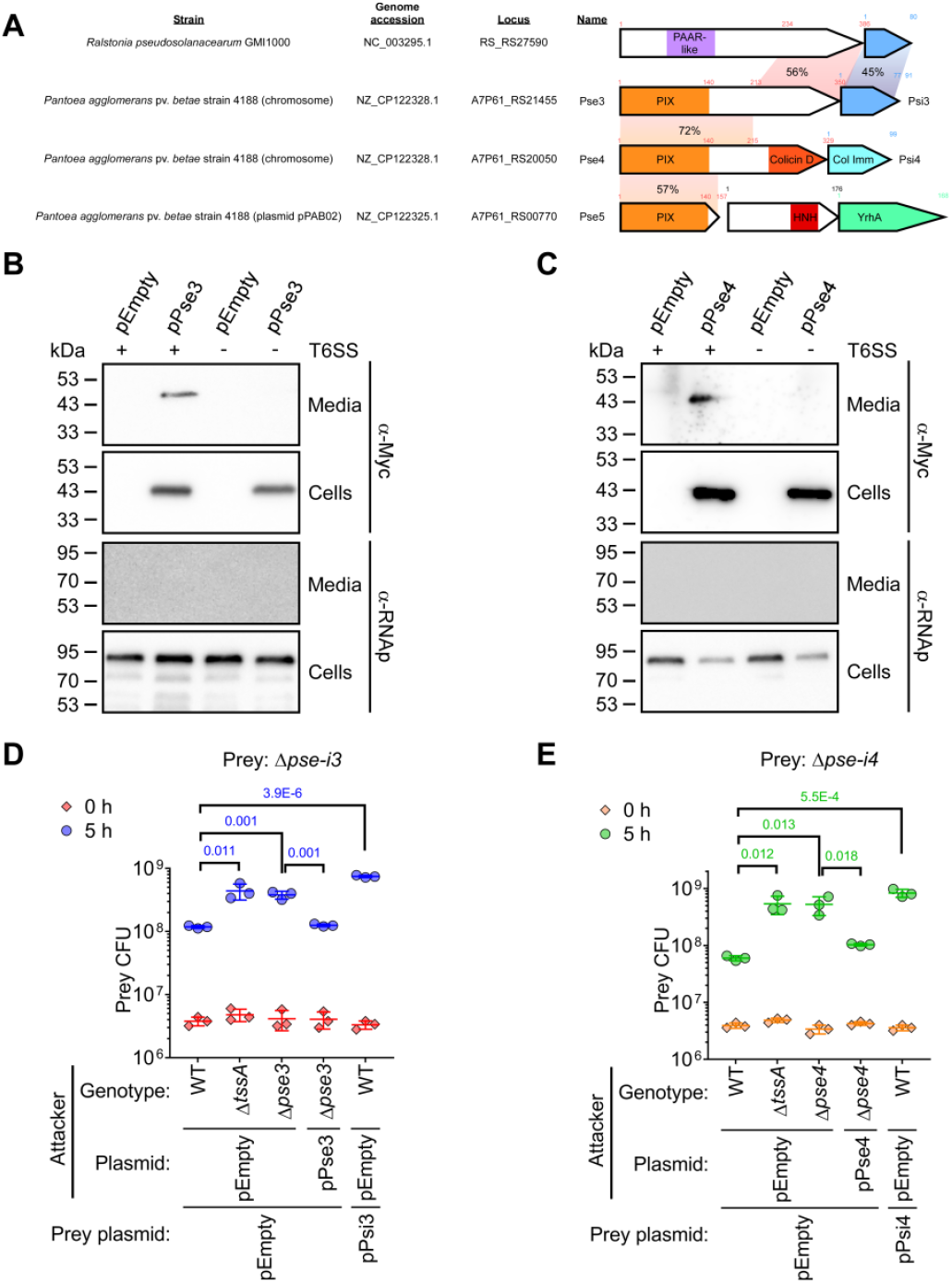
Pse3 and Pse4 are antibacterial T6SS effectors. **A)** The gene structure of the operons encoding WP_011003305.1, Pse3, Pse4, and Pse5. Arrows denote the direction of gene transcription; amino acid numbers are denoted above each gene; known and predicted domains are denoted inside the arrows. Colored rectangles denote regions of amino acid sequence homology; identity percentages are indicated. **B-C)** Expression (cells) and secretion (media) of C-terminally Myc-tagged Pse3 (B) and Pse4 (C) expressed from an arabinose-inducible plasmid in wild-type (WT) *Pab* or a T6SS^−^ mutant (Δ*tssA*). An empty plasmid (pEmpty) was used as a control. RNA polymerase sigma 70 (RNAp) was used as a loading and lysis control. **D-E)** Viability counts (CFU) of the indicated *Pab* prey strains carrying either an empty plasmid or a plasmid for the arabinose-inducible expression of Psi3 (pPsi3) or Psi4 (pPsi4) before (0 h) and after (5 h) co-incubation with the indicated *Pab* attacker strain carrying an empty plasmid or a plasmid for the arabinose-inducible expression of Pse3 (pPse3) or Pse4 (pPse4). The statistical significance between samples at the 5 h timepoint was calculated using an unpaired, two-tailed Student’s *t*-test. P values are denoted above. Data are shown as the mean ± SD; n = 3.

Pse3 and Pse4 are encoded upstream of small genes encoding WP_033759756.1 and WP_022625183.1, hereafter referred to as Psi3 and Psi4, respectively (**Fig. 1A**). Psi4 has a colicin immunity domain (CDD). Notably, Pse3 and Pse4 are not encoded near a known T6SS-associated gene. Based on these observations, we hypothesized that Pse3 and Pse4 are orphan antibacterial T6SS cargo effectors and that Psi3 and Psi4 are their cognate immunity proteins. To test this hypothesis, we first monitored the secretion of C-terminal Myc-tagged Pse3 and Pse4 expressed from a plasmid in *Pab*. As shown in **Fig. 1B**,**C**, both proteins were secreted to the medium from the wild-type strain but not from a mutant in which we inactivated the T6SS by deleting the conserved structural component, *tssA*. These results confirm that Pse3 and Pse4 are secreted by the *Pab* T6SS.

Next, we used bacterial competition assays to determine whether Pse/i3 and Pse/i4 are antibacterial T6SS effector and immunity pairs. Deletion of *pse3* and *psi3* (Δ*pse-i3*) rendered a prey strain sensitive to an attack by a wild-type strain (**Fig. 1D**). The toxicity was abolished when the attacker’s T6SS was inactivated (Δ*tssA*) and when *pse3* was deleted (Δ*pse3*), as well as upon expression of Psi3 from a plasmid (pPsi3) in the sensitive prey strain (**Fig. 1D**). Moreover, complementation of Pse3 from a plasmid (pPse3) restored the toxicity of a Δpse3 mutant. We obtained similar results for Pse4 and Psi4 (**Fig. 1E**). The growth of all *Pab* mutants was comparable to that of the wild-type strain (**Supplementary Fig. S2**). Taken together, these results indicate that Pse3 and Pse4 are antibacterial T6SS effectors and that Psi3 and Psi4 are their cognate immunity proteins.

### An N-terminal domain is necessary and sufficient for T6SS-mediated secretion of Pse3 and Pse4

AlphaFold2 structure predictions of Pse3 and Pse4 revealed two distinct domains in each protein (**Fig. 2A**,**B**) (Mirdita et al., 2022; Varadi et al., 2022). The effectors have a different predicted C-terminal domain toxic domain. Although the activity of the Pse3 C-terminal domain remains unknown, the C-terminal domain of Pse4 belongs to the Colicin_D nuclease superfamily. The predicted N-terminal domain is similar in both effectors (**Fig. 2C**). Since N-terminal domains found in modular T6SS effectors were previously shown to mediate effector delivery (Fridman et al., 2022; Jana et al., 2019; Kanarek et al., 2022; Pei et al., 2020), we hypothesized that the N-terminal domain found in Pse3 and Pse4 is a T6SS delivery domain. By comparing the sequence of Pse3 to its homologs, we identified a conserved region corresponding to amino acids 1-140 of Pse3 (**Fig. 2D**). This region corresponds with the N-terminal fold predicted by AlphaFold2 (**Fig. 2A,B** right panels and **Fig. 2C**); therefore, we named it the PIX (*<u>P</u>antoea* type s<u>IX</u>) domain.

**Figure 2.**
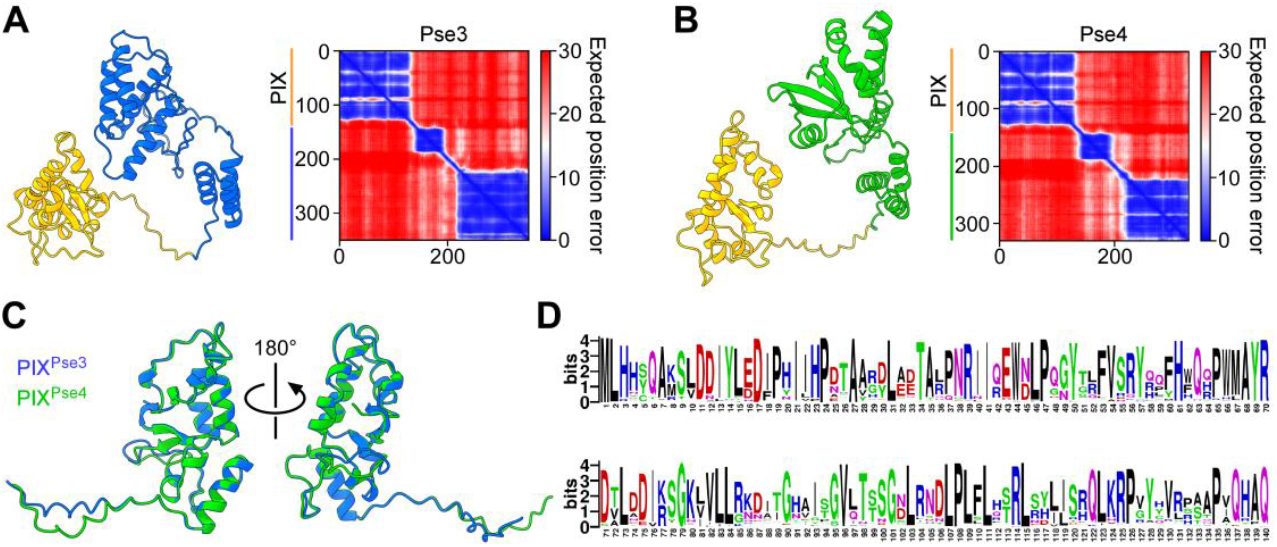
Pse3 and Pse4 contain a conserved N-terminal domain. **A-B)** AlphaFold2 structure predictions of Pse3 (A) and Pse4 (B). The predicted aligned error of each structure is shown on the right. A low predicted aligned error value indicates that the predicted relative position and orientation of the two residues are well-defined. Amino acids 1-140, corresponding to the PIX domain, are denoted in orange. **C)** Superimposition of the predicted PIX domain structures of Pse3 (blue) and Pse4 (green). **D)** The conservation logo of the PIX domain, based on multiple sequence alignment of sequences homologous to the N-terminus of Pse3. The position numbers correspond to the amino acids in Pse3.

Next, we sought to determine whether PIX is necessary and sufficient for T6SS-mediated secretion. Deletion of the first 37 amino acids of Pse3 and Pse4 (Pse3^Δ37^ and Pse4^Δ37^, respectively) abrogated their secretion from *Pab* (**Fig. 3A,B**), indicating that an intact PIX domain is necessary for effector secretion. Moreover, we found that the PIX domain, i.e., amino acids 1-140 of Pse3 and Pse4 (PIX^Pse3^ and PIX^Pse4^, respectively), is secreted from *Pab* in a T6SS-dependent manner, indicating that PIX is sufficient for T6SS-mediated secretion. Notably, the deletion of amino acids 141-200 (Δ141-200) in either effector did not affect their T6SS-mediated secretion, demonstrating that non-PIX domain regions that are homologous between the two effectors (**Fig. 1A**) are not required for T6SS-mediated secretion.

**Figure 3.**
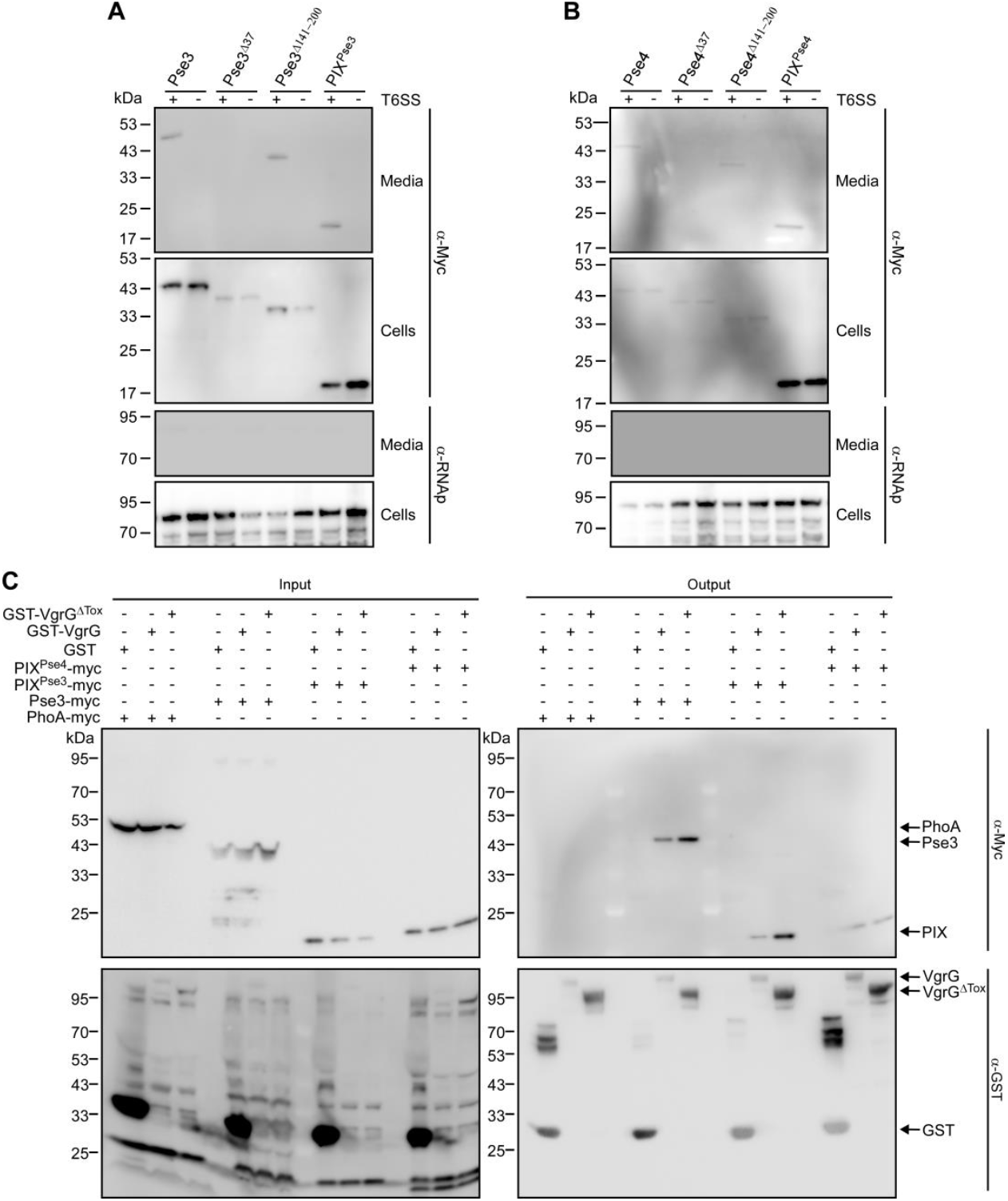
The PIX domain is necessary and sufficient for T6SS-mediated secretion and interaction with VgrG. **A-B)** Expression (cells) and secretion (media) of the indicated C-terminally Myc-tagged Pse3 (A) and Pse4 (B) forms expressed from an arabinose-inducible plasmid in wild-type (WT) *Pab* or a T6SS^−^ mutant (Δ*tssA*). RNA polymerase sigma 70 (RNAp) was used as a loading and lysis control. **C)** Glutathione beads-mediated pull-down analysis of the indicated GST fusion proteins mixed with lysates containing the indicated myc-tagged proteins. The *E. coli* PhoA lacking its N-terminal signal peptide was used as a negative control for non-specific interaction. The full-length VgrG used had an E752A mutation to inactivate its transglutaminase toxic domain. VgrG^ΔTox^, expressing amino acids 1-674 of VgrG.

Since various modular T6SS effectors with an N-terminal delivery domain interact with the secreted T6SS structural component VgrG (Ali et al., 2023; Jana et al., 2019; Pei et al., 2020), we hypothesized that Pse3 and Pse4 interact with the *Pab* VgrG via the PIX domain. Indeed, a pull-down assay revealed that the *Pab* VgrG (fused to GST, a glutathione S-transferase tag) interacts with the full-length Pse3, as well as with its PIX domain and with the PIX domain of Pse4, but not with the *Escherichia coli* PhoA protein used as a non-interacting negative control (**Fig. 3C**). Moreover, Pse3 and the PIX domains did not interact with GST alone, suggesting that the interaction between VgrG and the PIX domain is specific. We previously demonstrated that the *Pab* VgrG is a specialized antibacterial effector containing a C-terminal glucosaminidase toxic domain (Carobbi et al., 2022). We found that truncating this toxic domain does not hamper the interaction between VgrG and the PIX domain, indicating that PIX is not loaded onto the C-terminal toxic domain. This conclusion was further supported by a competition assay showing that a *Pab* strain in which we truncated the VgrG C-terminal toxic domain (Δ*vgrG*^*Tox*^) retains its ability to deliver Pse4 and intoxicate a Δ*pse-i4* prey strain (**Supplementary Fig. S3**). Taken together, our results reveal that the PIX domain, which is necessary and sufficient for T6SS-mediated secretion, is loaded onto the *Pab* specialized VgrG for T6SS-mediated delivery.

### The PIX domain defines a new class of orphan polymorphic T6SS effectors in *Enterobacterales*

The above results suggested that PIX is a T6SS delivery domain. As such, we reasoned that it could be used as a marker to identify T6SS effectors. To this end, we analyzed the distribution of PIX domains in bacteria. We found PIX at the N-termini of 224 unique proteins, encoded by 1571 bacterial genomes; 92.4% of these genomes harbor a T6SS gene cluster (i.e., at least 9 of 11 conserved T6SS core components were identified), further supporting the conclusion that PIX is a T6SS-associated delivery domain (**Supplementary Dataset S1**). PIX-encoding genomes predominantly belong to bacteria of the order *Enterobacterales*, including the genera *Citrobacter, Cronobacter, Enterobacter, Escherichia, Klebsiella, Pantoea, Salmonella*, and *Serratia* (**Supplementary Fig. S4A**). Remarkably, none of the identified PIX-encoding genes are found within a T6SS gene cluster or neighboring T6SS-associated genes, indicating the PIX-containing effectors are orphans (**Supplementary Dataset S1**). Moreover, analysis of the C-termini of PIX-containing proteins revealed several known toxic domains, such as NAD(P)^+^-degrading (NADase)-like, HNH and colicin-like nucleases, CdiA-CT-like, DUF2335 phospholipase, DUF3289, transglycosylase, and cysteine hydrolase-like (**Table 1** and **Supplementary Fig. S4B**). Interestingly, we could not assign a predicted activity to three types of C-terminal domains, suggesting that they represent novel toxins (i.e., Unknown_1/2/3 in **Table 1**). Most PIX-containing proteins (∼95%) are encoded immediately (i.e., within 25 bp) upstream of a gene that may encode a cognate immunity protein, suggesting that they mediate antibacterial activities.

**Table 1.** PIX-associated C-terminal extensions.

| Predicted C-terminal activity/domain <sup>a,b,c</sup> | Representative accession number | Number of unique proteins identified | An annotated putative immunity present <sup>d</sup> |
| --- | --- | --- | --- |
| NADase-like <sup>a</sup> | WP_223540050.1 | 74 | Yes |
| Unknown_1 | WP_064704874.1 (Pse3) | 64 | Yes |
| HNH nuclease <sup>b</sup> | WP_193405261.1 | 40 | Yes |
| Colicin D <sup>b</sup> | WP_022625183.1 (Pse4) | 20 | Yes |
| CdiA-CT <sup>a</sup> | WP_071198818.1 | 7 | Yes |
| DUF2335 (phospholipase) <sup>b</sup> | WP_017348567.1 | 3 | No<sup>*</sup> |
| DUF3289 <sup>b</sup> | WP_123225445.1 | 3 | Yes |
| Truncated DUF3289 <sup>b</sup> | WP_094107295.1, WP_118894928.1 | 3, 1 | - |
| Unknown_2 | WP_232486732.1 | 1 | Yes |
| Transglycosylase <sup>a</sup> | WP_027252378.1 | 1 | No<sup>*</sup> |
| Cysteine hydrolase-like <sup>a</sup> | WP_051640458.1 | 1 | Yes |
| Unknown_2 | WP_182808315.1 | 1 | Yes |
| Truncated HNH nuclease <sup>c</sup> | WP_250378716.1 | 2 | - |

<sup>a</sup> Predicted using HHpred, <sup>b</sup> Predicted using NCBI's Conserved Domain Database, <sup>c</sup> Predicted by homology to other proteins, <sup>d</sup> Assessed manually by inspecting the downstream encoded proteins, <sup>\*</sup> an unannotated open reading frame was manually identified; the table includes only proteins that have at least 60 amino acids C-terminal to the PIX domain

## Discussion

Identifying cargo effectors, especially orphan ones, remains one of the major challenges in the T6SS field. Here, we described a new N-terminal T6SS delivery domain, PIX, that defines a class of polymorphic toxins and can thus serve as a marker for new effectors. We used PIX to identify hundreds of PIX-containing proteins in T6SS-encoding *Enterobacterales* genomes, many of which have known or predicted C-terminal toxic domains. Since PIX-containing proteins are orphan (i.e., they do not neighbor known T6SS components in the genome), their identification as T6SS effectors would not have possible had it not been for PIX, at least not by using available computational approaches. Our findings underline the importance of annotating and investigating secretion system-associated delivery domains in polymorphic toxin classes, since they can be used as tools that enable novel effector identification. Indeed, we found three C-terminal domains fused to PIX with no known or predicted activity, suggesting that they are novel toxic domains.

We showed that PIX is necessary and sufficient for the T6SS-mediated secretion of polymorphic effectors through binding to VgrG. PIX is loaded onto a specialized VgrG containing a C-terminal toxic domain in *Pab*. However, PIX does not bind to the toxic domain. This finding indicates that T6SSs diversify the toxic payload that can be delivered in a single firing event by loading cargo effectors onto specialized effectors. A similar phenomenon was reported in *Vibrio cholerae*, where the cargo effector TleL binds the specialized effector VgrG3 (Dong et al., 2013).

PIX appears to be the first example of an order-specific T6SS delivery domain; we found it predominantly in *Enterobacterales*. Only a handful of other domains that define a class polymorphic T6SS cargo effectors are known: MIX, FIX, RIX, and Rhs repeats. Of these, only the distribution of the RIX domain is restricted to members of the *Vibrionaceae* family (Kanarek et al., 2022). This recently observed phenomenon of family- or order-specific T6SS delivery domains is intriguing. One possible explanation is that the PIX domain evolved in a common ancestor. However, since PIX-containing proteins are orphans and are often encoded next to DNA mobility elements, such as transposases, integrases, and prophages (**Supplementary Dataset S1**), they are probably shared via horizontal gene transfer that should not be restricted to this order. Another possibility is that PIX domains require an unknown protein present in *Enterobacterales* to fold or load onto the T6SS properly, or that they load onto a T6SS landing pad that is restricted to T6SSs found only in *Enterobacterales*. The mechanism that governs this restricted phylogenetic distribution of polymorphic T6SS effector classes remains to be investigated.

In conclusion, we identified a new class of orphan polymorphic T6SS effectors widespread in *Enterobacterales*. The PIX domain adds to a handful of known T6SS-specific effector delivery domains, and as such, it can reveal new insights into T6SS effector secretion and evolution.

## Materials and Methods

### Strains and media

For a complete list of strains used in this study, see **Supplementary Table S1**. *Escherichia coli* strains were grown in 2xYT (1.6% [wt/vol] tryptone, 1% [wt/vol] yeast extract, 0.5% [wt/vol] NaCl), lysogeny broth (LB, 1% [wt/vol] tryptone, 0.5% [wt/vol] yeast extract, and 0.5% [wt/vol] NaCl) or LB agar (1.5% [wt/vol]) plates at 37°C, or at 28°C when harboring effector-expressing plasmids. The media were supplemented with chloramphenicol (35 μg/mL), ampicillin (50 μg/mL), or kanamycin (50 μg/mL) to maintain plasmids, and with D-glucose (0.4% [wt/vol]) to repress expression from the P*bad* promoter. L-arabinose (0.1% to 0.05% [wt/vol]) or 0.1 mM IPTG were added to induce expression from the P*bad* or P*tac* promoters, respectively. *Pantoea agglomerans* pv. *betae* (*Pab*) and its derivatives were grown in LB or on LB agar supplemented with rifampicin (50 µg/mL) at 28°C. Kanamycin (50 μg/mL) or chloramphenicol (35 μg/mL) were added to the media to maintain plasmids when required.

### Plasmid construction

For a complete list of plasmids used in this study, see **Supplementary Table S2**. The coding sequences (CDS) of *pse3* (WP_064704874.1), *pse4* (WP_011003305.1), *psi3* (WP_033759756.1), *psi4* (WP_119918756.1) and derivative truncations lacking the first 37 amino acids *pse3*^*Δ37*^ and *pse4*^*Δ37*^ were PCR amplified from *Pab* genomic DNA. The CDS of *phoA* lacking its signal peptide (amino acids 1-21) was PCR amplified from *E. coli* BL21 (DE3). For the *pse3*^*Δ140-200*^ and *pse4*^*Δ140-200*^ constructs, the plasmids carrying the full length of *pse3* and *pse4* were used to amplify the corresponding sequences using primers designed adjacent to the deleted nucleotides and sharing complementarity at the 5’ end. The catalytically inactive version of *vgrG* and *vgrG*^*ΔTox*^ were amplified from previously reported pBAD/*Myc*-His constructs (Carobbi et al., 2022). The amplification products were inserted into the multiple cloning site (MCS) of pBAD/*Myc*-His or pBAD33.1, in-frame with a C-terminal *Myc*-His tag, or into pGEX4T-1 in-frame with an N-terminal GST tag using the Gibson assembly method (Gibson et al., 2009). The plasmids were transformed into *E. coli* via electroporation or into *Pab* via conjugation. Transconjugants were selected on LB agar plates supplemented with the appropriate antibiotics and 0.4% (wt/vol) D-glucose to repress expression from the P*bad* or P*tac* promoters.

### Construction of deletion strains

The construction of *Pab* mutant strains *ΔtssA* and Δ*vgrG*^*Tox*^ was described previously (Carobbi et al., 2022). For the in-frame deletion of *pse3, pse-i3, pse4* and *pse-i4* in *Pab*, 1 kb sequences upstream and downstream of each gene were subcloned into pDM4, a Cm^R^OriR6K suicide plasmid (O’Toole et al. 1996) and inserted into *E. coli* DH5α (λ-*pir*) by electroporation, and then into *Pab* via conjugation. Transconjugants were selected on LB agar plates supplemented with chloramphenicol (35 μg/mL), and then counter-selected on LB agar plates containing 15% (wt/vol) sucrose for loss of the sacB-containing plasmid. Deletions were confirmed by PCR.

### Secretion assays

*Pab* wild-type and *ΔtssA* strains containing arabinose-inducible expression plasmids for the expression of the indicated effector protein were grown overnight in LB media; the media were supplemented with kanamycin (50 μg/mL) to maintain the plasmids and 0.4% (wt/vol) D-glucose to repress expression from the P*bad* promoter. The cultures were washed and resuspended at a final OD600 of 0.15 in 5 mL of LB supplemented with kanamycin and 0.001% L-arabinose to induce protein expression from the plasmids. Bacterial cultures were incubated with constant shaking (220 rpm) at 28°C for 5 h. For expression fractions (cells), cells equivalent to 0.3 OD600 units were collected and cell pellets were resuspended in 40 µL of 2x Tris-glycine SDS sample buffer (Novex, Life Sciences). For secretion fractions (media), the remaining supernatant volumes were filtered (0.22 µm) and proteins were precipitated using the deoxycholate and trichloroacetic acid method (Bensadoun et al. 1976). Precipitated proteins were pelleted and washed twice with cold acetone, prior to re-suspension in 20 µL of 100 mM Tris-Cl (pH = 8.0), 20 µL of Tris-glycine SDS sample buffer (Novex, Life Sciences), and 1 μL of 1 N NaOH to maintain a basic pH. The expression and secretion samples were resolved on SDS-polyacrylamide gel electrophoresis (SDS-PAGE), transferred onto PVDF membranes using Trans-Blot Turbo Transfer (Bio-Rad), and immunoblotted with anti-*Myc* antibodies (Santa Cruz Biotechnology; 9E10, sc-40) or anti-RNA polymerase sigma 70 (RNAp) antibodies (BioLegend ; mouse mAb #663205) for loading and lysis control, at a 1:1000 dilution. Proteins were visualized using enhanced chemiluminescence (ECL). Results of a representative experiment out of at least three are shown.

### *Pab* growth assays

Triplicates of overnight-grown *Pab* cultures were normalized to OD600 = 0.01 in LB and transferred to a 96-well plate (200 μL per well). The 96-well plate was incubated in a microplate reader (BioTek SYNERGY H1) at 28°C with agitation (205 cpm), and growth was measured as OD600 at 10-minute intervals.

### Bacterial competition assays

Bacterial competition assays were performed as previously described (Carobbi et al., 2022), with minor changes. Briefly, the indicated attacker and prey strains were grown overnight in LB and then normalized to OD600 = 0.4. Attacker and prey cultures were mixed at a 4:1 (attacker:prey) ratio and the mixtures were spotted on LB agar plates supplemented with 0.05% (wt/vol) L-arabinose to induce protein expression from the arabinose-inducible plasmids. The plates were incubated at 28°C for 5h. The colony forming units (CFU) of the prey strain were determined by growing the mixtures from the 0 h and 5 h timepoints on selective plates. The assays were performed at least three times, and results of a representative experiment are shown.

### Pull-down assays

Overnight-grown cultures of *E. coli* BL21 (DE3) containing a plasmid for the arabinose-inducible expression of the indicated C-terminally *Myc*-tagged PhoA, Pse3 and Pse4 forms, or containing the IPTG-inducible plasmid for the expression of the indicated N-terminal GST-tagged VgrG forms or GST alone were diluted 1:20 in fresh 2xYT media supplemented with the appropriate antibiotics. The cultures were grown for 1.5 h at 37°C with agitation (215 rpm). Protein expression was induced by adding 0.05% (wt/vol) L-arabinose or 0.1 mM IPTG, followed by incubation at 28°C for 6 h with agitation (215 rpm). Cells were then harvested by centrifugation and kept at -80°C. The cell pellets were thawed and resuspended in lysis buffer (20 mM Tris-Cl pH 7.5, 500 mM NaCl, 5% [vol/vol] glycerol, 1 mM PMSF, 0.2 mg/mL lysozyme, and 20 µg/mL DNase). After incubation for 30 min at room temperature, cell debris were removed by centrifugation for 20 min at 17,000 x *g* at 4°C. The soluble protein fraction of cells expressing GST-tagged proteins was incubated with glutathione resin (GenScript; L00206) at 4°C for 1 h with constant rotation (10 rpm). After three washes with wash buffer (20 mM Tris-HCl pH 7.5, 200 mM NaCl, 5% [vol/vol] glycerol, and 1 mM EDTA), the resin was mixed with equal amounts of each bacterial lysate containing a *Myc*-tagged protein and with BSA at a final concentration of 0.25% (wt/vol). At this time, 50 μL of the mixed protein samples were taken and added to 50 μL of 2x Tris-glycine SDS sample buffer (Novex, Life Sciences). These samples were boiled at 95°C for 5 min and kept at -20°C to be used as the assay input fractions. The remaining samples were incubated overnight at 4°C with constant rotation (10 rpm). The resin was then collected by centrifugation at 370 x *g* at 4°C and washed twice with 5 mL of wash buffer. Bound proteins were eluted from the resin using 250 µL of ice-cold GST elution buffer (50 mM Tris-Cl pH 8 and 20 mM reduced glutathione), and 100 µL of the eluted fractions (output) were mixed with 100 µL of 2x Tris-glycine SDS sample buffer (Novex, Life Sciences) and boiled at 95°C for 5 min. Input and output fractions were resolved on SDS-PAGE and subjected to immunoblot analysis using anti-*Myc* antibodies (Santa Cruz Biotechnology; sc-40) or anti-GST (Santa Cruz Biotechnology; sc-459) antibodies at a 1:1000 dilution. Proteins were visualized using enhanced chemiluminescence (ECL). Results of a representative experiment out of at least three are shown.

### Protein structure prediction and visualization

Pse3 and Pse4 structures and their associated expected position error were predicted in ColabFold: AlphaFold2 using MMseqs2 (Mirdita et al., 2022). The PDB files used in this work are available as **Supplementary File S1**. Protein structures were visualized using ChimeraX 1.5 (Pettersen et al., 2021).

### Identifying PIX-containing proteins

The position-specific scoring matrix (PSSM) of PIX was constructed using the N-terminal 140 residues of Pse3 (WP_064704874.1) from *Pab*. Five iterations of PSI-BLAST (Altschul et al., 1997) were performed against the RefSeq protein database. In each iteration, a maximum of 500 hits with an expect value threshold of 10^−6^ were used. RPS-BLAST was then performed to identify PIX-containing proteins. The results were filtered using an expect value threshold of 10^−9^ and a minimal coverage of 70%. The genomic neighborhoods of PIX-containing proteins (**Supplementary Dataset S1**) were analyzed as described previously (Dar et al., 2018; Fridman et al., 2020). Duplicated protein accessions appearing in the same genome in more than one genomic accession were removed if the same downstream protein existed at the same distance. The T6SS core components in the PIX-containing bacterial genomes (**Supplementary Dataset S1**) were identified as previously described (Jana et al., 2019).

### Illustration of the conserved residues of the RIX domain

PIX domain sequences were aligned using Clustal Omega (Madeira et al., 2022). Aligned columns not found in the PIX domain of Pse3 were discarded. The PIX domain-conserved residues were illustrated using the WebLogo server (Crooks et al., 2004) (https://weblogo.berkeley.edu/logo.cgi).

### Constructing a phylogenetic tree of PIX-encoding bacterial strains

DNA sequences of *rpoB* were aligned using MAFFT v7 FFT-NS-i (https://mafft.cbrc.jp/alignment/server) (Katoh et al., 2002, 2018). The *rpoB* sequences were first clustered using CD-HIT to remove identical sequences (100% identity threshold). Partial and pseudogene sequences were discarded. The evolutionary history was inferred using the neighbor-joining method (Katoh et al., 2018) with the Jukes-Cantor substitution model (JC69). The analysis included 491 nucleotide sequences and 3,954 conserved sites.

### Analyses of PIX C-terminal sequences

The 100 C-terminal amino acids of PIX-containing proteins were clustered in two dimensions using CLANS (Frickey and Lupas, 2004). To predict the activities or domains in each cluster, at least two representative sequences (when more than one was available) were analyzed using the NCBI Conserved Domain Database (Marchler-Bauer et al., 2007) and HHpred (Gabler et al., 2020; Zimmermann et al., 2018). The presence or absence of a possible immunity gene immediately downstream of the PIX-encoding gene (i.e., within 50 base pairs of the PIX-encoding gene’s stop codon, as previously described (Fridman et al., 2020)) was determined.

## Data Availability

The experimental and computational data that support the findings of this research are available in this article and its supplementary information files, or upon request from the corresponding author.

## Acknowledgments

Sadly, Prof. Guido Sessa passed away shortly before the completion of this work. Prof. Sessa had a profound impact on the lives and scientific careers of all authors, and he will be sorely missed. We dedicate this manuscript to his memory, honoring his outstanding contributions to science. The work was supported by grants from the Israel Science Foundation (ISF; grant no. 488/19 to GS; grant no. 1362/21 to DS and EB). We thank members of the Sessa, Salomon, and Bosis laboratories for helpful discussions.

## Author contributions

Study conceptualization, DS and GS; funding acquisition, EB, DS, and GS; investigation and analysis, AC, SdN, EB, and DS; methodology, AC, EB, and DS; Writing – original draft, DS. All authors reviewed and approved the manuscript.

