## Supplemental Information for "An N-terminal delivery domain defines a new class of polymorphic T6SS effectors in *Enterobacterales*"

**Supplementary Figures S1-S4**

**Supplementary Tables S1-S2**

**Supplementary Dataset S1**

**Supplementary File S1**

**Supplementary References**

### Supplementary Figures

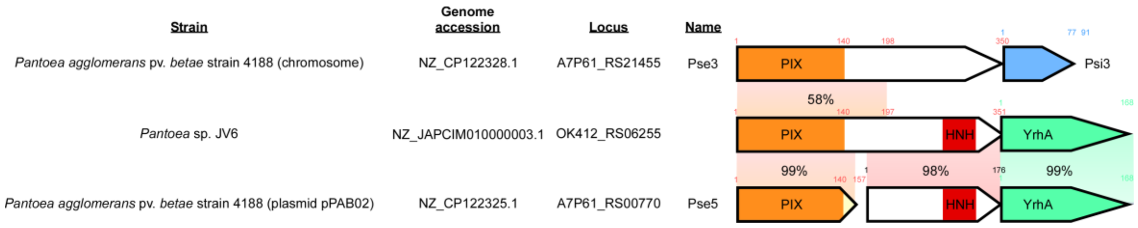


**Supplementary Figure S1. Pse5 is a putative truncated toxin.** The gene structure of the operons encoding Pse3, WP_089414745.1, and Pse5. Arrows denote the direction of gene transcription; amino acids numbers of note are denoted above each gene; known and predicted domains are denoted inside the arrows. Colored rectangles denote regions of amino acid sequence homology; identity percentages are indicated.

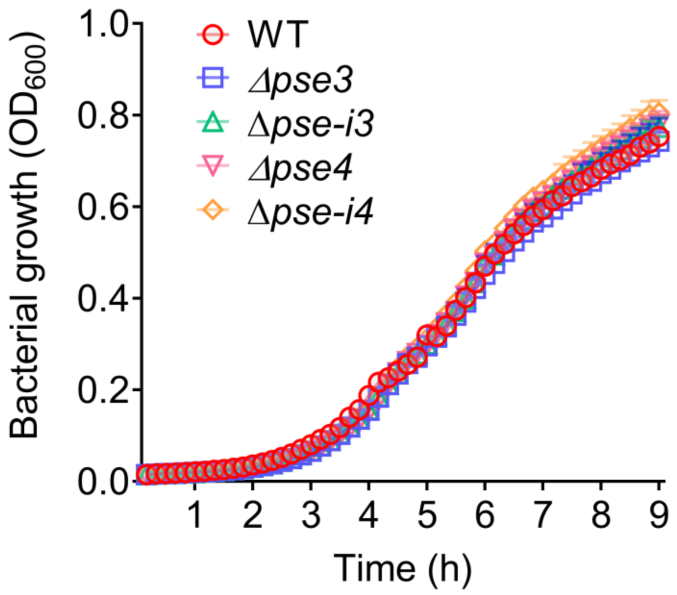


**Supplementary Figure S2. Deletion of *pse3* and *pse4* does not affect bacterial growth.** Growth of the indicated *Pab* strains, as determined by the optical density at 600 nm ($OD_{600}$). WT, wild-type. Data are shown as the mean ± SD; n = 3.

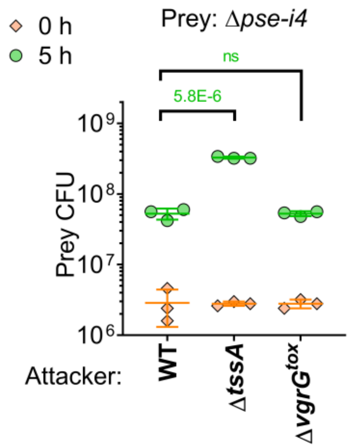


**Supplementary Figure S3. The C-terminal toxic domain of VgrG is not required for Pse4 delivery.** Viability counts (CFU) of the indicated *Pab* prey strain before (0 h) and after (5 h) co-incubation with the indicated *Pab* attacker strains. The statistical significance between samples at the 5 h timepoint was calculated using an unpaired, two-tailed Student's *t*-test. P values are denoted above. Data are shown as the mean ± SD; n = 3. Ns, no significant difference (P > 0.05).

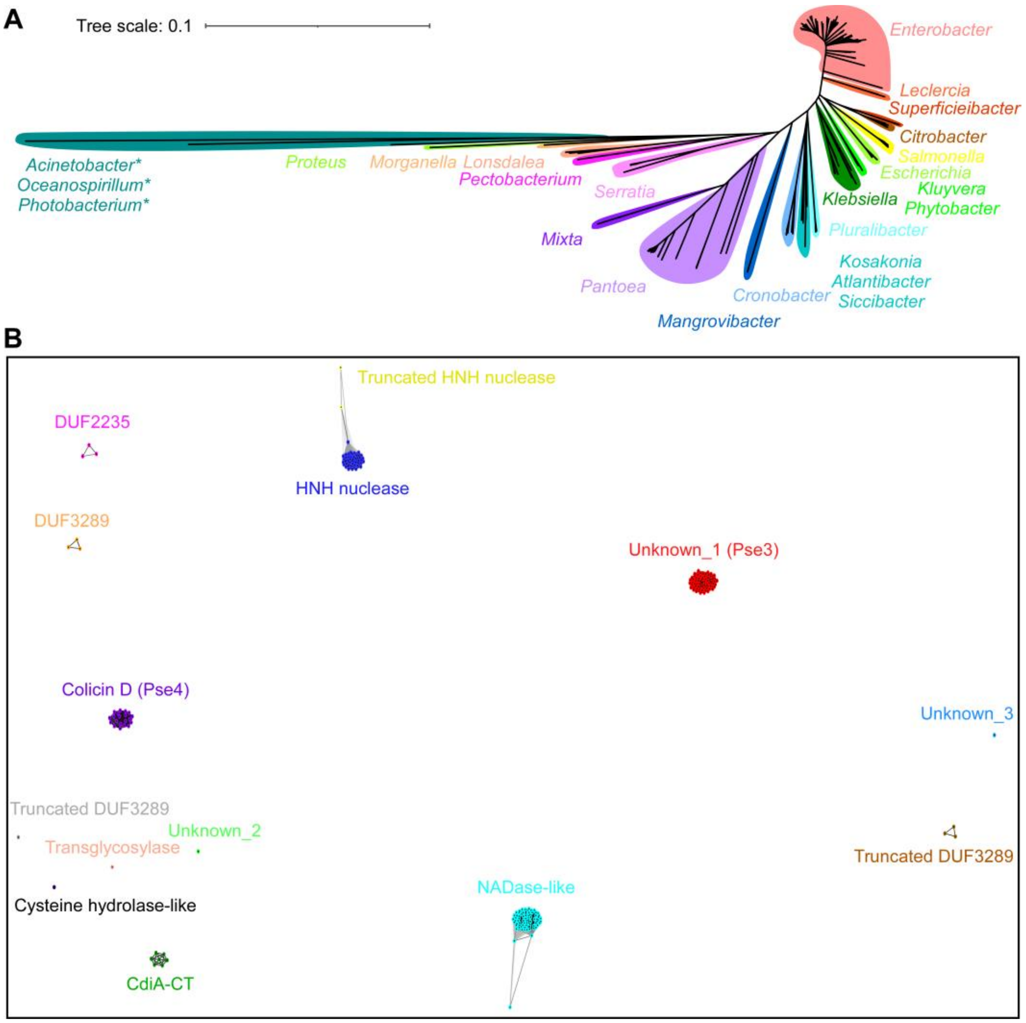


**Supplementary Figure S4. PIX-containing proteins are widespread in *Enterobacterales* and contain diverse predicted C-terminal toxic domains. A)** The phylogenetic distribution of bacteria encoding a protein with a PIX domain, based on the DNA sequence of *rpoB* coding for DNA-directed RNA polymerase subunit beta. The evolutionary history was inferred using the neighbor-joining method. Genera names are denoted. Asterisks denote bacterial genera that do not belong to the *Enterobacterales* order. **B)** C-termini (100 amino acids) of PIX-containing proteins were clustered in two dimensions based on all-against-all sequence similarity, with nodes representing unique sequences and connecting lines representing the distances between sequences. The predicted activities or domains identified in each cluster are denoted and are color-coded according to the nodes.

**Supplementary Tables**

**Supplementary Table S1. Bacterial strains used in this study.**

| Strain name | Genotype | Comments | Source |
|---|---|---|---|
| *Pantoea agglomerans* pv. *betae* 4188 (*Pab*) | Wild-type | Used as a template for PCR amplification, in secretion assays, and as an attacker in competition assays | Obtained from Isaac Barash (Valinsky et al., 1998) |
| *Pab* Δ*tssA* | Δ*tssA* | *Pab* derivative. Used in secretion assays and as an attacker in competition assays | (Carobbi et al., 2022) |
| *Pab* Δ*pse3* | Δ*pse3* | *Pab* derivative. Used in secretion assays and as an attacker in competition assays | This study |
| *Pab* Δ*pse4* | Δ*pse4* | *Pab* derivative. Used in secretion assays and as an attacker in competition assays | This study |
| *Pab* Δ*pse-i3* | Δ*pse3 and psi3* | *Pab* derivative. Used as prey in competition assays | This study |
| *Pab* Δ*pse-i4* | Δ*pse4 and psi4* | *Pab* derivative. Used as prey in competition assays | This study |
| *Pab* Δ$vgrG^{Tox}$ | Deletion of the C-terminal toxic domain of VgrG | *Pab* derivative. Used as an attacker in competition assays | Carobbi et al. 2022 |
| *Escherichia coli* DH5α | K-12 derivative laboratory strain | Used for cloning | Lab stocks |
| *Escherichia coli* DH5α (λ-*pir*) | K-12 derivative laboratory strain containing λ-*pir* | Used for plasmid maintenance and cloning | A gift from Eric V. Stabb |
| *Escherichia coli* BL21 (DE3) | laboratory strain | Used for protein expression in pull-down assays | Lab stocks |

**Supplementary Table S2. Plasmids used in this study.**

| Plasmid name | Description | Comments | Source |
|---|---|---|---|
| pDM4 | a $Cm^R$ and $oriV_{R6K}$-containing suicide vector | Used to generate deletions in the *Pab* genome | (O'Toole et al., 1996) |
| pDM4:*pse3* | pDM4 containing 1 kb upstream and 1 kb downstream of *pse3* in its MCS | Used to delete *pse3* from *Pab* | This study |
| pDM4:*pse-i3* | pDM4 containing 1 kb upstream *pse3* and 1 kb downstream of *psi3* in its MCS | Used to delete *pse3* and *psi3* from *Pab* | This study |
| pDM4:*pse4* | pDM4 containing 1 kb upstream and 1 kb downstream of *pse4* in its MCS | Used to delete *pse4* from *Pab* | This study |
| pDM4:*pse-i4* | pDM4 containing 1 kb upstream *pse4* and 1 kb downstream of *psi4* in its MCS | Used to delete *pse4* and *psi4* from *Pab* | This study |
| pDM4:*vgrG*$^{\Delta Tox}$ | pDM4 containing 1 kb upstream and 1 kb downstream of the *vgrG* C-terminal toxic domain (amino acids 674-829), in its MCS | Used to delete the C-terminal toxic domain of *vgrG* from *Pab* | (Carobbi et al., 2022) |
| pBAD/*Myc*-His | Arabinose-inducible pBAD/*Myc*-His plasmid harboring a $Kan^R$ cassette | Used for cloning and arabinose-inducible expression | (Salomon et al., 2013) |
| pBAD/*Myc*-His:*pse3* | pBAD/*Myc*-His containing the *Pab* effector *pse3*; the gene is cloned in-frame with the C-terminal *Myc*-His tag of the plasmid | Used for Pse3 inducible expression in secretion assays and in pull-down | This study |
| pBAD/*Myc*-His:*pse3*$^{\Delta 141\text{-}200}$ | pBAD/*Myc*-His containing the *pse3* lacking the codons corresponding to amino acids 141-200; the gene is cloned in-frame with the C-terminal *Myc*-His tag of the plasmid | Used for Pse3$^{\Delta 141\text{-}200}$ inducible expression in secretion assays | This study |
| pBAD/*Myc*-His:*pse3*$^{\Delta 37}$ | pBAD/*Myc*-His containing the *pse3* lacking the codons corresponding to amino acids 1-37; the gene is cloned in-frame with the | Used for Pse3$^{\Delta 37}$ inducible expression in secretion assays | This study |

| | | | |
|---|---|---|---|
| | C-terminal *Myc*-His tag of the plasmid | | |
| pBAD/*Myc*-His:*pse3*$^{1\text{-}140}$ | pBAD/*Myc*-His containing the *Pab* effector *pse3* codons corresponding to amino acids 1-140; the gene is cloned in-frame with the C-terminal *Myc*-His tag of the plasmid | Used for Pse3$^{1\text{-}140}$ inducible expression in secretion assays and in pull-down | This study |
| pBAD/Myc-His:*pse4* | pBAD/*Myc*-His containing the *Pab* effector *pse4*; the gene is cloned in-frame with the C-terminal *Myc*-His tag of the plasmid | Used for Pse4 inducible expression in secretion assays and in pull-down | This study |
| pBAD/*Myc*-His:*pse4*$^{\Delta 141\text{-}200}$ | pBAD/*Myc*-His containing the *pse4* lacking the codons corresponding to amino acids 141-200; the gene is cloned in-frame with the C-terminal *Myc*-His tag of the plasmid | Used for Pse4$^{\Delta 141\text{-}200}$ inducible expression in secretion assays | This study |
| pBAD/*Myc*-His:*pse4*$^{\Delta 37}$ | pBAD/*Myc*-His containing the *pse4* lacking the codons corresponding to amino acids 1-37; the gene is cloned in-frame with the C-terminal *Myc*-His tag of the plasmid | Used for Pse4$^{\Delta 37}$ inducible expression in secretion assays | This study |
| pBAD/*Myc*-His:*pse4*$^{1\text{-}140}$ | pBAD/*Myc*-His containing the *Pab* effector *pse4* codons corresponding to amino acids 1-140; the gene is cloned in-frame with the C-terminal *Myc*-His tag of the plasmid | Used for Pse4$^{1\text{-}140}$ inducible expression in secretion assays and in pull-down | This study |
| pBAD:*phoA* | pBAD/*Myc*-His containing the *E. coli* K-12 alkaline-phosphatase *phoA* lacking the codons encoding amino acids 1-21; the gene is cloned in-frame with the C-terminal *Myc*-His tag of the plasmid | Used for PhoA expression in pull-down assays | This study |
| pGEX4T-1 | Lactose analog isopropyl β-D-thiogalactoside (IPTG)-inducible pGEX plasmid harbouring an Amp$^R$ cassette | Used as negative control bait in pull-down assays | GE Healthcare |

| | | | |
|---|---|---|---|
| pGEX4T-1:*vgrG*$^{E752A}$ | pGEX4T-1 containing the *Pab vgrG* gene carrying an E752 substitution | Used as bait in pull-down assays | This study |
| pGEX4T-1:*vgrG*$^{\Delta Tox}$ | pGEX4T-1 containing the *Pab vgrG* gene corresponding to amino acids 1-645 | Used as bait in pull-down assays | This study |
| pBAD33.1 | pBAD series expression plasmid encoding the $Cm^R$ gene and the p15A origin of replication | Used for selective grow of *Pab* prey strains during self-competiton assays | Addgene |
| pBAD33.1:*psi3* | pBAD33.1 containing the *Pab psi3* gene; the gene is cloned in-frame with a C-terminal *Myc*-His tag | Used for Psi3 expression in competition assays | This study |
| pBAD33.1:*psi4* | pBAD33.1 containing the *Pab psi4* gene; the gene is cloned in-frame with a C-terminal *Myc*-His tag | Used for Psi4 expression in competition assays | This study |

### Supplementary Datasets

**Supplementary Dataset S1. PIX containing homologs, their genomic neighborhoods, and the presence of T6SS in their encoding genomes.**

### Supplementary Files

**Supplementary File S1. PDB files of Pse3 and Pse4 structures predicted by AlphaFold2.**
